# FiNNpy 2.0: Fast MEG source reconstruction

**DOI:** 10.1101/2024.10.11.617925

**Authors:** David Crompton, Prerana Keerthi, Luka Milosevic, Maximilian Scherer

## Abstract

Herein, we present the 2.0 update of FiNNpy, which expands the toolkit’s initial scope from the analysis of multi-site electrophysiological activity to track information propagation to include source reconstruction capability for MEG signals. Following the toolkit’s design guidelines, the new functionality has been optimized towards minimal resource consumption, making source reconstruction much faster, especially when executed in a parallelized fashion.

Furthermore, several quality-of-life aspects are introduced to support source reconstruction, such as grouping source-space activity into cortical areas, a cohesive documentation & in-code extensive literature references, and wrapping of previously terminal exclusive functions of FreeSurfer.

## I. INTRODUCTION

Source reconstruction is a crucial step in magnetoencephalographic (MEG) signal processing. The purpose is twofold: to project noninvasively acquired electrophysiological activity onto the cortical surface, and, with additional steps, into a shared space for group analyses. As parameters (e.g. head orientation and shape) vary across recordings/subjects, source projections have to be computed frequently. While approaches solving this mathematically underdetermined problem may be computed through various methods, efficient implementations thereof have received little attention.

With the 2.0 update, FiNNpy, a Python-based toolbox for the analysis of electrophysiological signals [1], provides time- and memory-efficient routines for source reconstruction. As these have been streamlined for parallelized execution (including external tools), signals may be projected into source-space significantly faster. Several quality-of-life improvements have been added to the reconstruction pipeline, to decrease workload during initial setup and execution. To support curious users in advancing their expertise in this domain, the processing pipeline’s individual steps have been thoroughly documented, including references to theoretical aspects of the source reconstruction effort.

Originally, FiNNpy had been developed to investigate multi-site information propagation, to *Fi*nd *N*europhysiological *N*etworks. Therefore, the toolbox implements a wide range of methods to quantify these phenomena. For phase amplitude coupling (PAC), FiNNpy implements the modulation index [2], phase lock value [3], mean vector length [4], and the direct modulation index [5]. For spectral coherence, the toolbox implements magnitude squared coherence [6], imaginary coherence [7], weighted phase lag index [8], phase slope index [9], and directional absolute coherence [10].

This functionality has now been expanded with the MEG source reconstruction pipeline presented herein. Improvements to speed, resource consumption, and precision were quantified in comparison to a contemporary alternative. With the 2.0 release, FiNNpy is available via pip with the command “pip install finnpy” and available for direct download on GitHub (https://github.com/neurophysiological-analysis/FiNN).

## II. METHODS

### A. Data

Data from https://openneuro.org/ was used [11], namely *sub-Z446*.

### B. Assessing speed and memory consumption

The source reconstruction execution time and memory consumption were compared between FiNNpy and MNE [12], another Python-based toolbox offering similar functionality. To maintain comparability, evaluations were performed on the same hardware (CPU: AMD Ryzeon 9 5950X; RAM: 64 GB; OS: Arch Linux). Of note, all evaluations were performed without parallelization and with multithreading flags disabled as parallelization benefits herein are primarily bound by the ratio of RAM required to the amount available, as well as the number of unoccupied threads. For example, if a single source reconstruction effort requires 25 % of available RAM, execution of 4 source reconstruction efforts concurrently can provide an up to a 4 times increase in speed, if *≤* 4 unoccupied threads are available. Execution speed and RAM consumption for each assessment pipeline were measured repeatedly across 4 runs. Time expired was quantified as the nanoseconds between the end and start of an assessment pipeline. RAM consumption as the difference in system-wide RAM occupancy between prior (5 to 10 s) to and during pipeline execution. Memory consumption was tracked indirectly as some processing steps involving third-party tools spawn dedicated subprocesses, making direct RAM evaluations difficult. Both metrics were normalized with respect to MNE’s average time/memory consumption as a point of reference.

Separate pipelines were created for FiNNpy and MNE, both calculating a sensor-to-source space projection matrix from the output of the FreeSurfer pipeline for anatomical surface extraction. Performance was quantified across the entire pipelines for both FiNNpy and MNE. As FiNNpy implements noise normalization via dynamic statistical parameter mapping (dSPM; [13]), this approach was chosen for both pipelines.

### C. Precision of eigenvalue/-vector decompositions

Eigenvalue decompositions are employed to co-register MRI & MEG orientations/positions and to estimate sensor channels’ covariance. As these computations require the execution of multiple operations beyond the precision supported by 64 bit floats, numerical instability may arise, leading to a decrease in the accuracy of the sensor-to-source space projection matrix. To quantify this effect, eigenvalue decompositions required in the FiNNpy & MNE pipelines were computed with 16, 32, 64(default in Python), 128, and 256 bit precision. Differences in accuracy were quantified by calculating the Euclidean distance between the eigenvectors of 16, 32, 64, 128 bit versus the eigenvectors of 256 bit. Results were weighted by the eigenvalues of the 256 bit computation. To derive a single error score, the error scores for each precision were summed.

### D. Code

The full evaluation code is available on GitHub at: https://github.com/VoodooCode14/FiNNpy2.0eval. The framework is available on GitHub at: https://github.com/neurophysiological-analysis/FiNN/tree/develop/finnpy.

## III. RESULTS

### A. FiNNpy 2.0 vs. MNE

Comparing single-core CPU performance, sensor-to-source space projection calculation post anatomy extraction (see above) was approximately 2 times faster, and required 2 times less memory (average & peak consumption) in FiNNpy over MNE (Figure 1).

**Fig 1:**
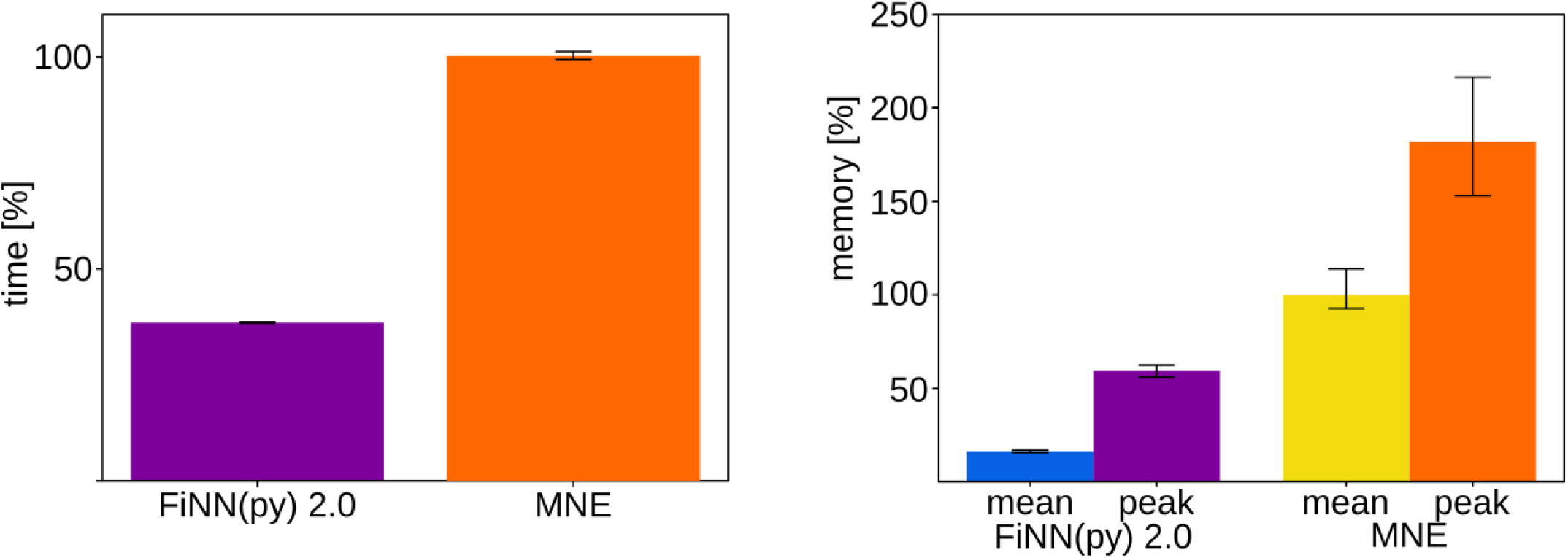
Single core processing time (left) and memory consumption (right) for MEG source reconstruction. Bars indicate mean values over 4 runs. Minimum and maximum values are indicated.

### B. Eigenvalue precision

Increasing numerical precision for eigenvalue decompositions resulted in an increase of precision. For example, comparing 128 bit and 64 bit eigenvalue decompositions, the lower precision resulted in a 13 % increase in error (metric: eigenvalue-weighted average Euclidean distance between eigen-vectors; Supplementary Figure 2).

## IV. DISCUSSION

Through source reconstruction, recordings of cortical activity acquired in sensor space may be projected onto the cortical surface. This provides an estimate of spatial origin. Calculating the sensor-to-source space transformation matrix is computationally complex. Furthermore, since key factors, such as head-to-sensor positioning/orientation and anatomy (e.g., thickness of the temporal lobe) vary between recordings and subjects, a dedicated transformation matrix is required for each case.

Herein, the authors present the 2.0 update of FiNNpy, providing efficient source reconstruction routines, optimized for speed and accuracy, supplemented with important quality-of-life improvements.

### A. Reconstruction speed

In FiNNpy 2.0, time required for source reconstruction is reduced via 2 approaches: First, by decreasing computational complexity in the reconstruction pipeline. Second, by simplifying the parallelized processing of multiple subjects. For the 1st approach, any source reconstruction steps following FreeSurfer’s MRI extraction have been optimized for efficient processing and minimal memory consumption. This reduced time and memory requirements by a factor of 2 and 3 respectively, compared to other contemporary Python-based solutions, allowing for up to 6 times faster execution if subjects are processed in parallel. For the 2nd approach, previously terminal-only calls of FreeSurfer have been simplified by providing Python-based wrappers. This enables corresponding functionality to be directly invoked from within Python, removing the need to switch between interfaces & programming languages. Furthermore, wrappers have been designed to be compatible with FiNNpy’s easy-to-use thread pool, or any other package that provides similar functionality.

### B. Reconstruction accuracy & numerical stability

Millimeter-scale errors in the sensor-to-source space projection matrix may cause cortical activity to be mapped onto an incorrect cortical area. However, the transformation’s accuracy is degraded by numerical inaccuracies (rounding errors). These arise from computations being performed across a wider range of values than supported by floating-point precision, magnified if done repeatedly.

FiNNpy 2.0 approaches this issue by identifying critical elements in the reconstruction routine wherein accumulating rounding errors may have a considerable impact on the reconstruction’s precision. Eigenvalue calculations were identified as critical elements, as these require a high number of calculations in their computation, herein performed across a wide range of numerical values. In the source reconstruction effort, eigenvalues and eigenvectors are used to determine MEG-to-anatomy orientations and the covariance between individual MEG channels, both important factors. Their computation was re-implemented in pure Python and Python-wrapped C++, up to 256 bit precision (from 64 bit). While the pure Python version is easier to use, the C++ one is much faster.

### C. Usability increases

Source reconstruction commonly scatters cortical activity onto thousands of cortical source points, significantly impairing any statistical effort on the group level. To address this, FiNNpy offers the ability to merge cortical source points into specific cortical areas as defined in the Desikan-Killiany atlas [14]. By merging activity from many cortical source points into fewer cortical areas, their spatial distribution is effectively smoothed. This homogenizes effects across subjects (in a spatially limited area), simplifying single-subject & group-level inferential analyses. Furthermore, by reducing the number of source points/areas by two orders of magnitude, the number of potential hypotheses in exploratory statistical analyses is greatly reduced. Importantly, the aforementioned homogenization and the smaller number of hypotheses reduce the statistical power required to detect statistically significant effects.

### D. Limitations

The source reconstruction effort was exemplary evaluated using a single MRI/MEG data set. While reconstruction time may vary between different MRI/MEG data sets, to some degree, the presented improvements in processing speed and resource consumption should be mostly unaffected.

## V. CONCLUSION

FiNNpy is a rapidly evolving toolbox, primarily aimed at neuroscientists with a background in data analytics seeking to apply complex methodology. Herein, the authors present FiNNpy’s 2.0 update, providing a MEG source reconstruction effort. Following the framework’s design philosophy to optimize functionality for minimal resource consumption, FiNNpy offers a rapid source-reconstruction routine supplemented by important quality-of-life improvements.

## REFERENCES

[1] M. Scherer, T. Wang, R. Guggenberger, L. Milosevic, and A. Gharabaghi, “FiNN: A toolbox for neurophysiological network analysis,” Network Neuroscience, pp. 1–34, Jul. 2022.

[2] A. B. L. Tort, R. Komorowski, H. Eichenbaum, and N. Kopell, “Measuring Phase-Amplitude Coupling Between Neuronal Oscillations of Different Frequencies,” Journal of Neurophysiology, vol. 104, no. 2, pp. 1195–1210, May 2010.

[3] F. Mormann, J. Fell, N. Axmacher, et al., “Phase/amplitude reset and theta-gamma interaction in the human medial temporal lobe during a continuous word recognition memory task,” Hippocampus, vol. 15, no. 7, pp. 890–900, 2005.

[4] R. T. Canolty, E. Edwards, S. S. Dalal, et al., “High Gamma Power Is Phase-Locked to Theta Oscillations in Human Neocortex,” Science, vol. 313, no. 5793, pp. 1626–1628, Sep. 2006.

[5] M. Scherer, T. Wang, R. Guggenberger, L. Milosevic, and A. Gharabaghi, “Direct modulation index: A measure of phase amplitude coupling for neurophysiology data,” Human Brain Mapping, Dec. 2022.

[6] R. Srinivasan, P. L. Nunez, and R. B. Silberstein, “Spatial filtering and neocortical dynamics: Estimates of EEG coherence,” IEEE transactions on bio-medical engineering, vol. 45, no. 7, pp. 814–826, Jul. 1998.

[7] G. Nolte, O. Bai, L. Wheaton, Z. Mari, S. Vorbach, and M. Hallett, “Identifying true brain interaction from EEG data using the imaginary part of coherency,” Clinical Neurophysiology, vol. 115, no. 10, pp. 2292–2307, Oct. 2004.

[8] M. Vinck, R. Oostenveld, M. van Wingerden, F. Battaglia, and C. M. A. Pennartz, “An improved index of phase-synchronization for electrophysiological data in the presence of volume-conduction, noise and sample-size bias,” NeuroImage, vol. 55, no. 4, pp. 1548– 1565, Apr. 2011.

[9] G. Nolte, A. Ziehe, V. V. Nikulin, et al., “Robustly Estimating the Flow Direction of Information in Complex Physical Systems,” Phys. Rev. Lett., vol. 100, no. 23, p. 234 101, Jun. 2008.

[10] M. Scherer, T. Wang, R. Guggenberger, L. Milosevic, and A. Gharabaghi, “Directional Absolute Coherence: A phase-based measure of effective connectivity for neurophysiology data,” p. 2022.02.07.479359, Feb. 2022.

[11] K. A. Fadeev, Ilacai V. Romero Reyes, Dzerassa D. Goiaeva, et al., Perception of vowel sounds in children with autism spectrum disorders and typically developing children (MEG/ERF study), Type: dataset, 2024.

[12] A. Gramfort, M. Luessi, E. Larson, et al., “MEG and EEG data analysis with MNE-Python,” Frontiers in Neuroscience, vol. 7, 2013.

[13] A. M. Dale, A. K. Liu, B. R. Fischl, et al., “Dynamic Statistical Parametric Mapping: Combining fMRI and MEG for High-Resolution Imaging of Cortical Activity,” Neuron, vol. 26, no. 1, pp. 55–67, Apr. 2000.

[14] R. S. Desikan, F. Sgonne, B. Fischl, et al., “An automated labeling system for subdividing the human cerebral cortex on MRI scans into gyral based regions of interest,” NeuroImage, vol. 31, no. 3, pp. 968–980, Jul. 2006.

