## Supplementary Material for "FiNNpy 2.0: Fast MEG source reconstruction"

David Crompton[1,2], Prerana Keerthi[1,2], Luka Milosevic[1,2,3], Maximilian Scherer[1,4]
Krembil Research Institute, University Health Network, Toronto, Canada. [1]
Institute of Biomedical Engineering, University of Toronto, Toronto, Canada. [2]
Center for Advancing Neurotechnological Innovation to Application (CRANIA), Toronto, Canada. [3]
Department of Neurology, University Hospital, LMU Munich, Munich, Germany [4]

### I. Additional results

#### *A. Accuracy comparison for eigenvector & eigenvalue computations*

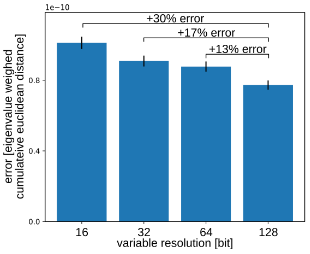


Fig. 1: Changes in eigenvector & eigenvalue accuracy for different levels of floating point precision.

As expected, increasing the precision of the floating-point variables used during eigenvalue & eigenvector calculations enhances the accuracy of the overall results.

Of note, the change in accuracy from $32\,\mathrm{bit}$ to $64\,\mathrm{bit}$ is roughly similar to the change between $64\,\mathrm{bit}$ and $128\,\mathrm{bit}$.
